# Operating regimes of covalent modification cycles at high enzyme concentrations

**DOI:** 10.1101/167825

**Authors:** Ronny Straube

## Abstract

The Goldbeter-Koshland model has been a paradigm for ultrasensitivity in biological networks for more than 30 years. Despite its simplicity the validity of this model is restricted to conditions when the substrate is in excess over the converter enzymes − a condition that is easy to satisfy *in vitro*, but which is rarely satisfied *in vivo*. Here, we analyze the Goldbeter-Koshland model by means of the total quasi-steady state approximation which yields a comprehensive classification of the steady state operating regimes under conditions when the enzyme concentrations are comparable to or larger than that of the substrate. Where possible we derive simple expressions characterizing the input-output behavior of the system. Our analysis suggests that enhanced sensitivity occurs if the concentration of at least one of the converter enzymes is smaller (but not necessarily much smaller) than that of the substrate and if that enzyme is saturated. Conversely, if both enzymes are saturated and at least one of the enzyme concentrations exceeds that of the substrate the system exhibits concentration robustness with respect to changes in that enzyme concentration. Also, depending on the enzyme’s saturation degrees and the ratio between their maximal reaction rates the total fraction of phosphorylated substrate may increase, decrease or change nonmonotonically as a function of the total substrate concentration. The latter finding may aid the interpretation of experiments involving genetic manipulations of enzyme and substrate abundances.

## 1. Introduction

The quasi-steady state approximation (QSSA) or its close relative, the rapid equilibrium approximation, are frequently used to derive reduced models for enzyme-catalyzed reaction networks (Segel and Slemrod, 1989; Straube et al., 2005; Salazar and Höfer, 2009; Radulescu et al., 2012). However, while this procedure mostly preserves the steady state structure of the network it often fails to correctly capture its transient dynamics. For this purpose the total QSSA (or tQSSA), which is based on certain linear combinations of the original variables, has proven to yield much better approximations, especially when the enzyme concentration becomes comparable to or larger than that of the substrate (Borghans et al., 1996; Tzafriri, 2003; Tzafriri and Edelman, 2004; Ciliberto et al., 2007). Here, we wish to show that the tQSSA can also be useful to classify the steady state behavior of a reaction network.

To this end, we consider the Goldbeter-Koshland model for covalent modification cycles (Fig. 1) which has been a paradigm for the generation of ultrasensitivity in signaling networks (Goldbeter and Koshland Jr., 1981; Tyson et al., 2003; Ferrell Jr. and Ha, 2014). Due to its simplicity it has also been used as a toy model to analyze generic network properties such as signal transmission (Tănase-Nicola et al., 2006; Tostevin and ten Wolde, 2009) or cross talk (Behar et al., 2007; Rowland et al., 2012). However, the applicability of the Goldbeter-Koshland model is restricted to conditions when the substrate concentration is much higher than that of the converter enzymes. While this condition is routinely used to conduct *in vitro* experiments it is rarely satisfied *in vivo* (Blüthgen et al., 2006; Legewie et al., 2008). To overcome this limitation we reanalyze the Goldbeter-Koshland model by means of the tQSSA. The resulting quadratic equations for the concentrations of the enzyme-substrate complexes are solved in the limit of high and low affinity which defines 4 operating regimes. Further analysis within each regime defines several subregimes the number of which depends on the enzyme affinities. Together, this yields a comprehensive classification of the steady state operating regimes for covalent modification cycles which extends the classification given by Gomez-Uribe et al. (Gomez-Uribe et al., 2007) to conditions when the enyzme concentrations are comparable to or larger than that of the substrate.

**Figure 1:**
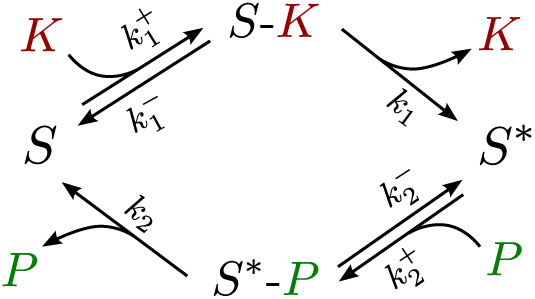
Reaction scheme for the Goldbeter-Koshland model describing phosphorylation/dephosphorylation cycles as mediated by a kinase (*K*) and a phosphatase (*P*). *S* and *S*\* denote the unphosphorylated and phosphorylated form of the substrate, respectively.

## 2. Material and Methods

### 2.1. QSSA vs. total QSSA

Assuming mass-action kinetics the dynamics of the network depicted in Fig. 1 is described by the ODE system

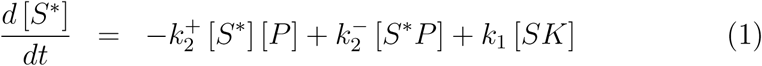

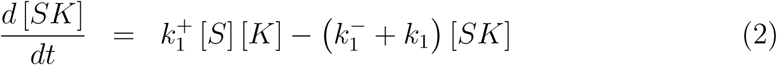

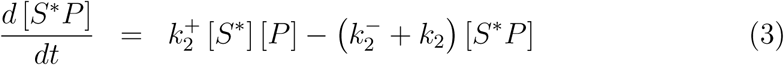

together with the conservation relations

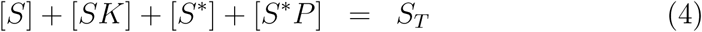

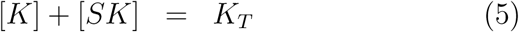

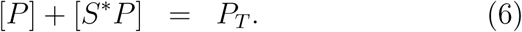

Here, *S_T_*, *K_T_* and *P_T_* denote total concentrations of substrate, kinase and phosphatase, respectively. Solving Eqs. (4) - (6) for [*S*], [*K*] and [*P*], and substituting the resulting expressions into Eqs. (1) - (3) yields a set of ODEs for [*S*\*], [*SK*] and [*SP*].

The QSSA relies on the assumption that the enzyme substrate complexes, *SK* and *S*P*, rapidly approach a quasi-steady state defined by

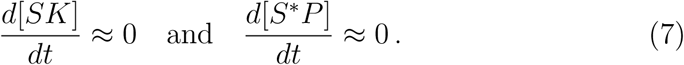

Solving these equations for [*SK*] and [*S*P*] yields the algebraic relations

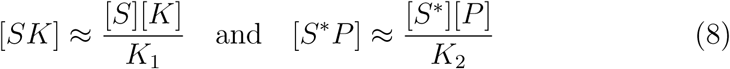

where 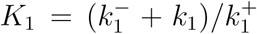 and 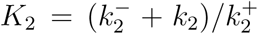 denote the Michaelis-Menten constants of the kinase and the phosphatase, respectively. Under the additional assumption that the substrate is in excess over the enzymes, i.e.

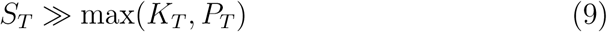

Goldbeter and Koshland derived the following ODE for [*S*\*] (Goldbeter and Koshland Jr., 1981)

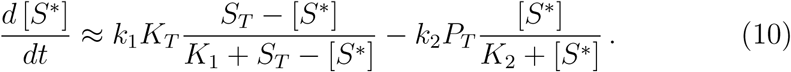

Note that under the substrate excess condition (Eq. 9) the conservation relation for the substrate (Eq. 4) simplifies to [*S*] + [*S*\*]≈ *S_T_*, i.e. the Goldbeter-Koshland model neglects sequestration of substrate into enzyme-substrate complexes.

One of the hallmarks of the Goldbeter-Koshland model is that it predicts ultrasensitive responses if both converter enzymes operate in saturation (Fig. 2A). In general, the steady state response curve (*d*[*S*\*]=*dt* = 0) is given by

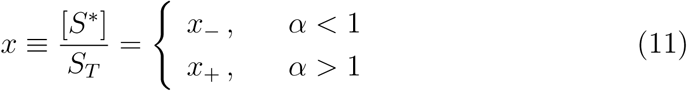

where *α* = *k*_1_*K_T_*/(*k*_2_*P_T_*) denotes the ratio between the maximal reaction rates of kinase and *x*_±_ denotes

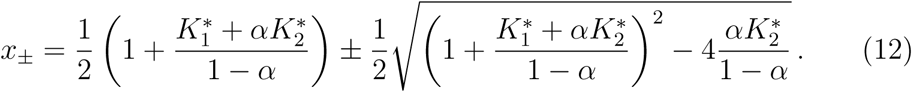

Hence, the steady state response within the Goldbeter-Koshland model can be classified by the magnitude of the rescaled Michaelis-Menten constants 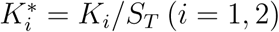. Apart from the ultrasensitive regime (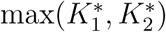 ≪ 1) there are three further steady state operating regimes denoted by (Gomez-Uribe et al., 2007): hyperbolic 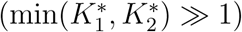, signal-transducing 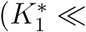 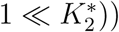 and threshold-hyperbolic 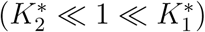. Comparing the predictions of the Goldbeter-Koshland model with those of full numerical solutions (cf. Fig. 2) we see that unless both enzymes are unsaturated (Fig. 2B) the predictions based on Eq. (12) overestimate the concentration of the phosphorylated substrate at large values of as sequestration effects are neglected.

**Figure 2:**
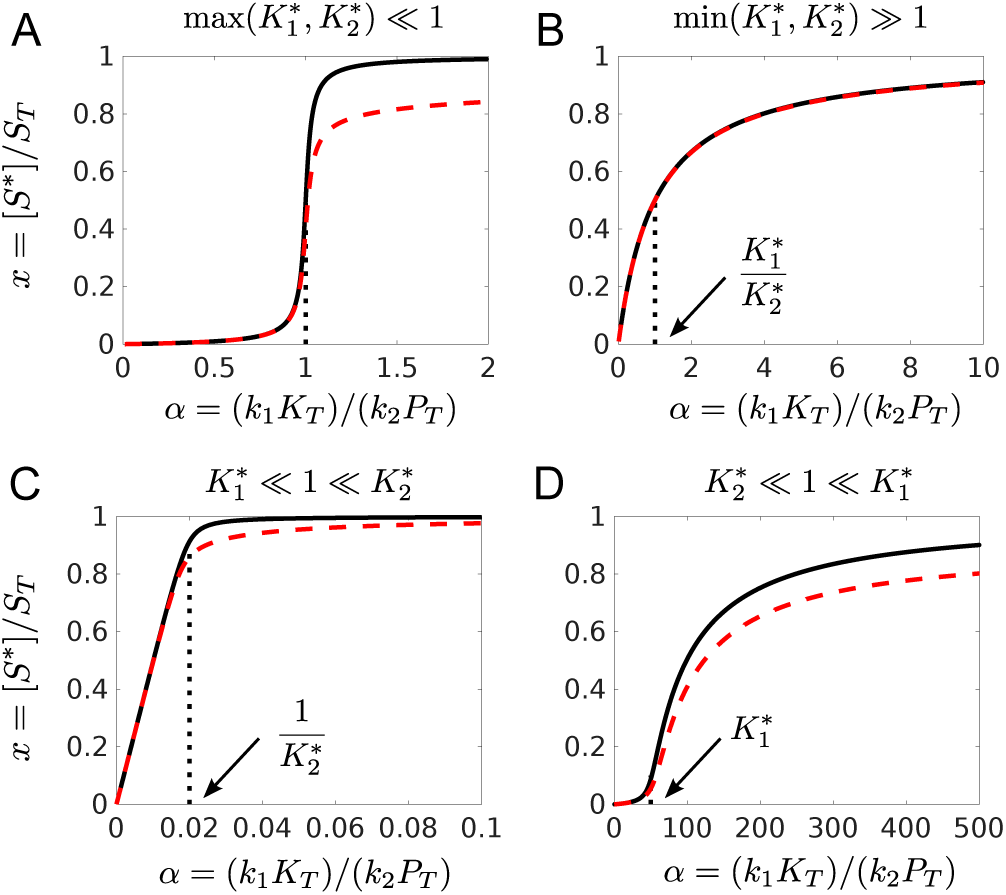
Steady state operating regimes of the Goldbeter-Koshland model under conditions of substrate excess: (A) Ultrasensitivity 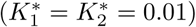, (B) hyperbolic response 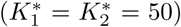, (C) signal-transducing response 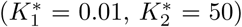, (D) threshold-hyperbolic response 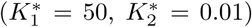. Solid lines were computed from Eq. (11). Dashed lines correspond to the numerical steady state solution of Eqs. (1) - (6). Dotted lines mark the parameter value for half-maximal activation (A,B) or the location of thresholds (C,D). Parameters: *K_T_* = *P_T_* = 0.1*μM*, *S_T_* = 1*μM*, 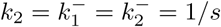.

The total QSSA starts by introducing the total concentrations of phosphorylated and unphosphorylated substrate as

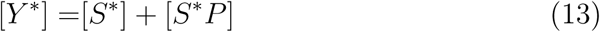

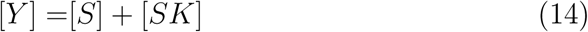

together with the inverse relations

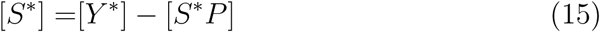

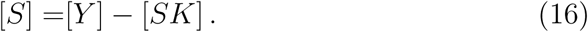

In the new variables, *Y*\* and *Y*, the conservation relation for the substrate (4) simplifies to

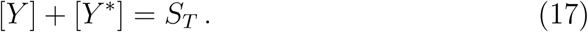

Addition of Eqs. (1) and (3) yields an ODE for [*Y*\*] which replaces that for [*S*\*] according to

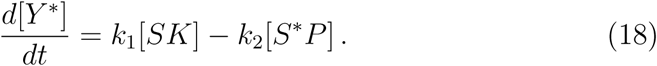

In the total QSSA the concentrations of the enzyme-substrate complexes are, again, determined by the QSSA relations (8), but now [*S*\*] and [*S*] have to be replaced according to Eqs. (15) and (16), respectively. In conjunction with the conservation relations (5), (6) and (17) this yields two quadratic equations for [*SK*] and [*S*P*] given by

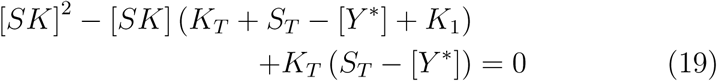

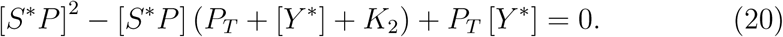

Taking into account the constraints 0 ≤ [*SK*] ≤ *K_T_* and 0 ≤ [*S*P*] ≤ *P_T_*, the solutions of Eqs. (19) and (20) are given by

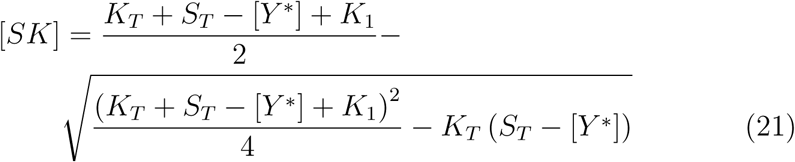

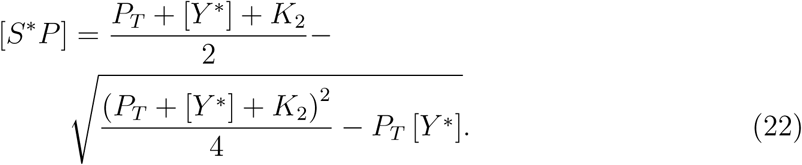

Substituting these expressions into Eq. (18) yields the tQSSA for the ODE system defined by Eqs. (1) - (6) and for initial conditions in the range 0 ≤ [*Y*\*] ≤ *S_T_*.

### 2.2. Operating regimes within the total QSSA

The steady state behavior within the Goldbeter-Koshland model can be classified by comparing the Michaelis-Menten constants with the total substrate concentration. However, the structure of the quadratic equations (19) and (20) suggests a different classification scheme within the total QSSA. In fact, quadratic equations of the form

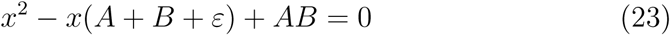

have been analyzed previously in the context of bifunctional enzymes (Straube, 2013) and two-component systems (Straube, 2014) as well as in the analysis of substrate competition (Buchler and Louis, 2008; Straube, 2015). Depending on the relative magnitude between *ε* and *A* (or *B*) the solution Eq. (23) can be approximated by

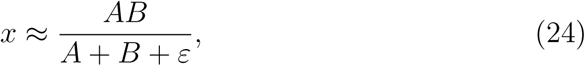

when *ε* ≫ min(*A, B*) or

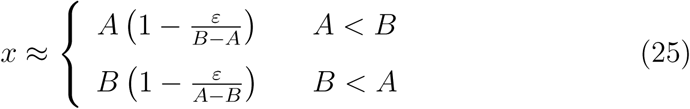

when *ε* ≪ max(*A, B*).

Applied to the quadratic equations (19) and (20) this suggests to classify the steady state behavior within the total QSSA by comparing the Michaelis-Menten constants with the enzyme concentrations (rather than the substrate concentration). In the high-*K_M_* regime, defined by *K*_1_ ≫ *K_T_* and *K*_2_ ≫ *P_T_*, this yields the approximate solutions

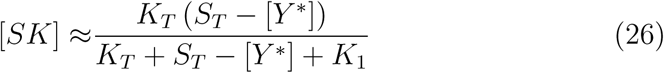

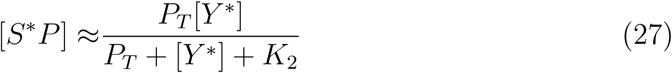

whereas in the low-*K_M_* regime (*K*_1_ ≪ *K_T_* and *K*_2_ ≪ *P_T_*) the approximations read

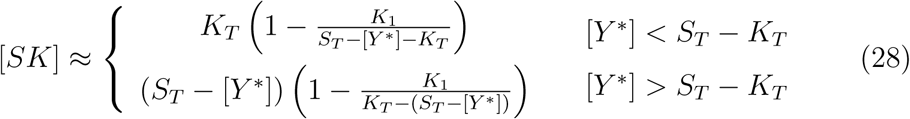

and

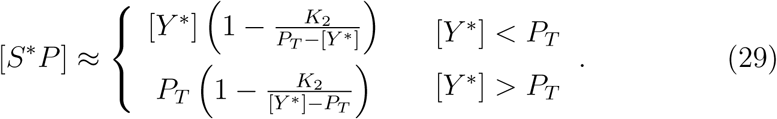

While previous studies only focussed on the high-*K_M_* regime, defined by Eqs. (26) and (27) (Borghans et al., 1996; Ciliberto et al., 2007; Gomez-Uribe et al., 2007), we shall also consider the low-*K_M_* regime as well as the mixed regimes.

## 3. Results

### 3.1. The low-K_M_ regime

To derive an explicit expression for [*Y*\*] in the low-*K_M_* regime (*K*_1_ ≪ *K_T_*and *K*_2_ ≪ *P_T_*) we need to combine the expressions in Eqs. (28) and (29) in an appropriate manner. To this end, we distinguish two cases:

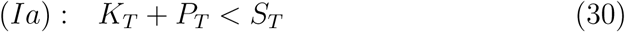

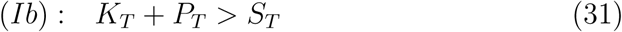

Note that in the first case [*Y*\*] < *P_T_* implies [*Y*\*] < *S_T_* − *K_T_* and vice versa in the second case. Hence, in the regime (Ia) we obtain (to lowest order) the tQSSA approximation

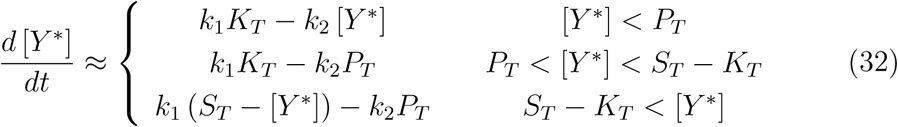

whereas in the second regime (Ib) the approximation reads

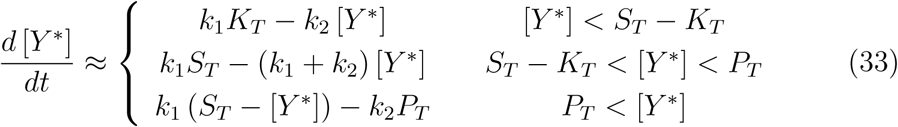

### 3.1.1. Response curve with respect to k_1_

*Regime (Ia): K_T_* + *P_T_ < S_T_*. To derive the steady state response curve as a function of *k*_1_ we set the first line in Eq. (32) to zero

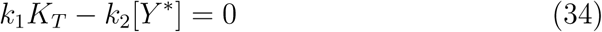

which has the solution [*Y*\*] = (*k*_1_/*k*_2_)*K_T_*. Applying the condition [*Y*\*] < *P_T_* yields *k*_1_ < *k*_2_*P_T_/K_T_* In the middle interval (second line in Eq. 32) there is no steady state solution to lowest order (unless *k*_1_*K_T_* = *k*_2_*P_T_*). To find the steady state solution in this interval one would have to consider the first-order correction terms for [*SK*] and [*S*P*] in Eqs. (28) and (29). However, for the present purpose this will not be necessary. Instead, we go ahead setting the last line in Eq. (32) to zero

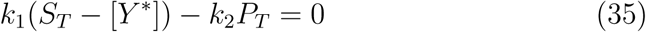

which has the solution [*Y*\*] = *S_T_* − (*k*_2_/*k*_1_)*P_T_*. Applying the condition *S_T_ − K_T_* < [*Y*\*] yields *k*_1_ > *k*_2_*P_T_/K_T_*. Combining these results shows that

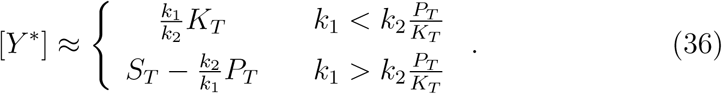

Hence, in the lowest order approximation the middle interval in Eq. (32) shrinks to zero. In fact, in a small interval around the threshold value *k*_1*c*_ = *k*_2_*P_T_/K_T_* the response curve changes in an ultrasensitive manner from (*k*_1_/*k*_2_)*K_T_* to *S_T_* − (*k*_2_/*k*_1_)*P_T_* (Fig. 3A) where the threshold (defined by *α* = 1) is the same as in the Goldbeter-Koshland model (cf. Eq. 11).

To compute the stimulus-response curve for the phosphorylated substrate we employ Eqs. (15) and (29) which yields

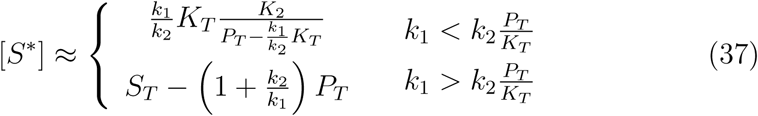

Hence, below *k*_1*c*_ the phosphorylation level is low (since *K*_2_/*P_T_* ≪ 1) whereas beyond the threshold it increases hyperbolically. Note that the steep increase of the response curve near *k*_1*c*_ is not captured by this approximation. However, from Eq. (37) we see that even at high kinase activity (*k*_1_ ≫ *k*_2_) the maximal phosphorylation level is not *S_T_*, but *S_T_ − P_T_*, which results from the sequestration of substrate by the phosphatase. Consistently, as the phosphatase concentration increases the maximal phosphorylation level drops, and the ultrasensitivity of the response curve (measured by the Hill coefficient *n_H_*) decreases (Fig. 3B). Note that even when *K_T_* + *P_T_ > S_T_* the system retains enhanced sensitivity.

*Regime (Ib): K_T_* + *P_T_ > S_T_*. The steady state equations resulting from the first and the third line in Eq. (33) are the same as those in Eq. (32). Applying the conditions [*Y*\*] < *S_T_ − K_T_* and *P_T_* < [*Y*\*] to the solutions [*Y*\*] = (*k*_1_/*k*_2_)*K_T_* and [*Y*\*] = *S_T_* − (*k*_2_/*k*_1_)*P_T_* yields *k*_1_ < *k*_2_(*S_T_ − K_T_*)/*K_T_* and *k*_1_ > *k*_2_*P_T_*/(*S_T_ − P_T_*), respectively. The steady state solution corresponding to the middle regime in Eq. (33) reads [*Y*\*] = *k*_1_*S_T_*/(*k*_1_ + *k*_2_). Applying the conditions [*Y*\*] > *S_T_ − K_T_* and [*Y*\*] < *P_T_* yields *k*_1_ > *k*_2_(*S_T_ − K_T_*)/*K_T_* and *k*_1_ < *k*_2_*P_T_*/(*S_T_ −P_T_*), respectively.

Combining these results shows that for *K_T_* + *P_T_ > S_T_* the response curve can be approximated by (cf. Fig. 3C)

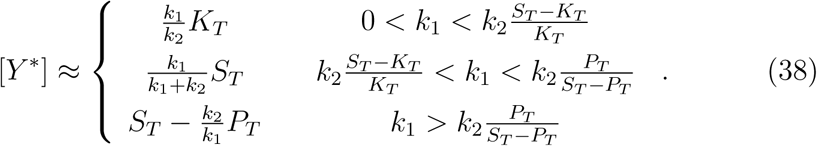

Hence, if *K_T_ < S_T_* and *P_T_ < S_T_* (with *K_T_* +*P_T_ > S_T_*) there are three distinct intervals in which [*Y*\*] increases with *k*_1_ in a specific manner. Since *k*_1_ has to be positive one or two of these intervals become empty when *K_T_ > S_T_* or*P_T_ > S_T_* in which case the solution simplifies. For example, if *K_T_ > S_T_* the first interval is empty and Eq. (38) becomes (cf. Fig. 3D)

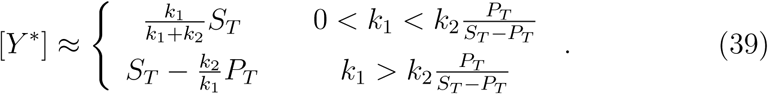

Similarly, if *P_T_ > S_T_* the solution simplifies to (cf. Fig. 3E)

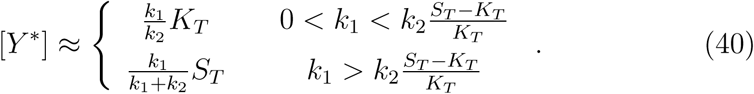

Finally, if both enzyme concentrations are larger than that of the substrate Eq. (38) becomes (cf. Fig. 3F)

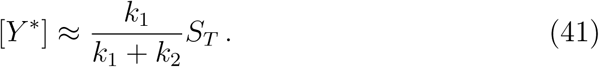

Together, this suggests that the system retains enhanced sensitivity (*n_H_* > 1) in the regime *K_T_* + *P_T_ > S_T_* if the concentration of at least one of the converter enzymes remains lower than that of the substrate.

From the expression in Eq. (41) we see that if the concentrations of both converter enzymes independently exceed the substrate concentration the Hill coefficient for the response curve becomes *n_H_* = 1 so that ultrasensitivity is lost. Interestingly, the shape of the stimulus-response curve does not depent on the enzyme concentrations in that regime. This suggests that the system exhibits concentration robustness (Shinar and Feinberg, 2010) with respect to *K_T_* and *P_T_*, i.e. changes in either concentration would not affect the steady state output of the system. Also, from the expressions in Eqs. (39) and (40) we see that the former does not depend on *K_T_* while the latter does not depend on *P_T_* suggesting that concentration robustness also exists if only one of the enzyme concentrations exceeds the substrate concentration.

Taken together, these results suggest that under the conditions *K_T_ <S_T_ < P_T_* and *P_T_ < S_T_ < K_T_* the system exhibits both ultrasensitivity and concentration robustness.

### 3.1.2. Response curve with respect to S_T_

Next we wish to analyze how the total fraction of phosphorylated substrate changes with the total substrate concentration. To this end, we rewrite Eq. (38) in terms of *S_T_*. As a first step we determine the boundaries of the intervals in which the three solution branches exist. From the first and the third line we see that the solutions [*Y*\*] = (*k*_1_/*k*_2_)*K_T_* and [*Y*\*] = *S_T_* − (*k*_2_/*k*_1_)*P_T_* exist for *S_T_* > (1 + *k*_1_/*k*_2_)*K_T_* and *S_T_* > (1 + *k*_2_/*k*_1_)*P_T_*, respectively. The solution in the middle interval ([*Y*\*] = *k*_1_*S_T_*/(*k*_1_ + *k*_2_)) exists for *S_T_* < min[(1 + *k*_1_/*k*_2_)*K_T_*, (1 + *k*_2_/*k*_1_)*P_T_*]. Hence, for *k*_1_*K_T_ < k*_2_*P_T_* (*α* < 1) the stimulus-response curve can be approximated by

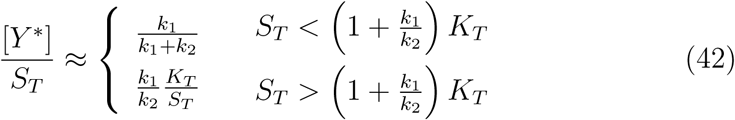

whereas for *k*_1_*K_T_ > k*_2_*P_T_* (*α* > 1) the approximation reads

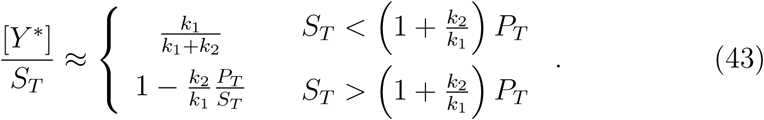

From these expressions it is apparent that the fraction of phosphorylated substrate may decrease (Eq. 42) or increase (Eq. 43) as a function of the total substrate concentration depending on the ratio *α* = *k*_1_*K_T_*/(*k*_2_*P_T_*) betweenthe maximal reaction rates of kinase and phosphatase (Fig. 4A).

**Figure 3:**
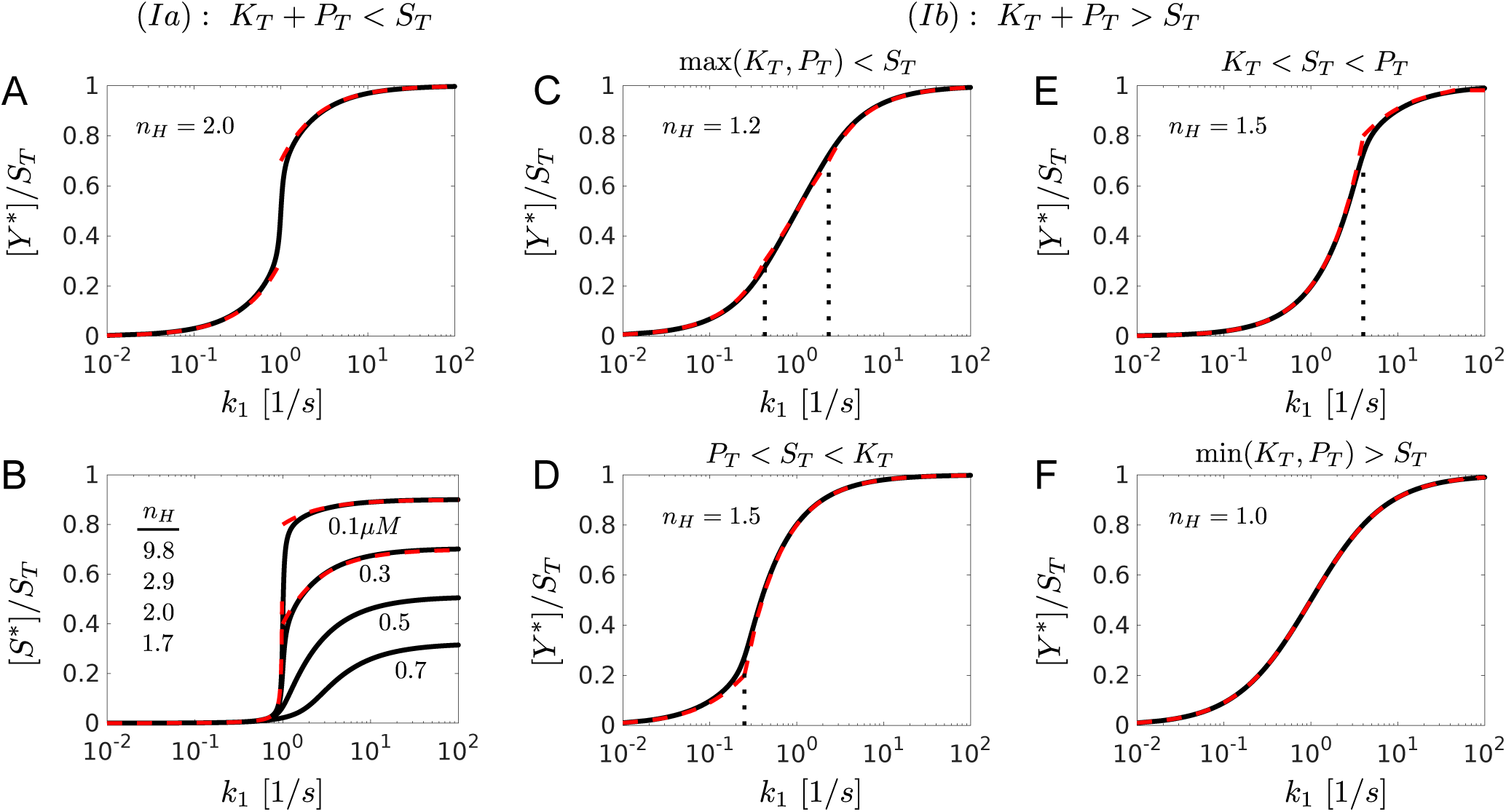
Response curves in the low-*K_M_* regime as a function of *k*_1_: (A) *K_T_* = *P_T_* = 0.3*μM*. (B) As the enzyme concentrations increase the maximal phosphorylation level as well as the ultrasensitivity decrease. Numbers indicate the value of *K_T_* = *P_T_* for *S_T_* = 1*μM*. The Hill coefficient is defined by *n_H_* = ln 81 = ln *R_S_* where 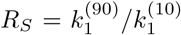 denotes the ratio of parameter values that elicite 90% (and 10%) of the maximal response. (C) *K_T_* = *P_T_* = 0.7*μM*. (D) *K_T_* = 2*μM*, *P_T_* = 0.2*μM*. (E) *K_T_* = 0.2*μM*, *P_T_* = 2*μM*.(F) *K_T_* = *P_T_* = 2*μM*. Black solid lines correspond to the numerical steady state solution of Eqs. (1) - (6). Red dashed lines were computed from Eq. (36) (A), Eq. (37) (B), Eq. 38 (C), Eq. 39 (D), Eq. 40 (E), Eq. 41 (F). Other parameters: *S_T_* = 1*μM*, *K*_1_ = *K*_2_ = 0.01*μM*, *k*_2_ = 1/*s*.

**Figure 4:**
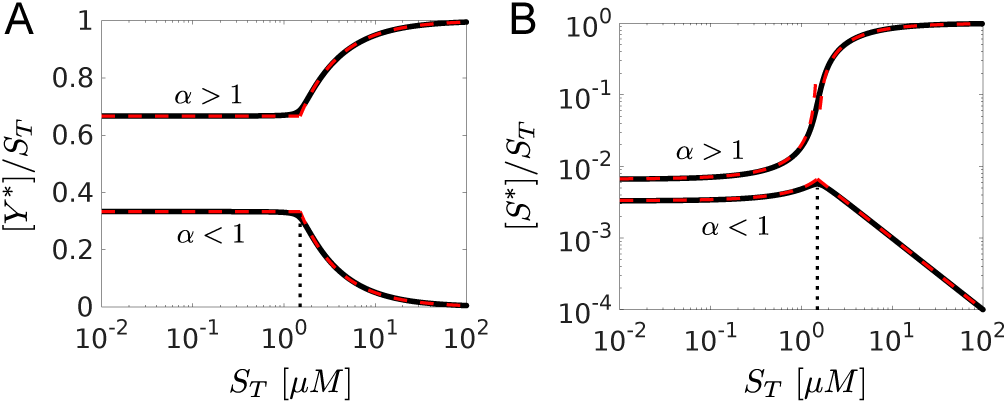
Response curves in the low-*K_M_* regime as a function of *S_T_* for different values of *α* = *k*_1_*K_T_*/(*k*_2_*P_T_*): (A) total fraction of phosphorylated substrate. (B) fraction of phosphorylated substrate. Black solid lines correspond to the numerical steady state solution of Eqs. (1) - (6). Red dashed lines were computed from Eqs. (42) - (45). Vertical dotted lines correspond to the thresholds (1 + *k*_1_/*k*_2_)*K_T_* (upper curves) and (1 + *k*_2_/*k*_1_)*P_T_* (lower curves). Parameters: *k*_2_ = 0.5/*s* (upper curves), *k*_2_ = 2/*s* (lower curves). Other parameters: *k*_1_ = 1/*s*, *K*_1_ = *K*_2_ = 0.01*μM*, *K_T_* = *P_T_* = 1*μM*.

To compute approximate expressions for [*S*\*]/*S_T_* we employ Eqs. (15) and (29) which yields

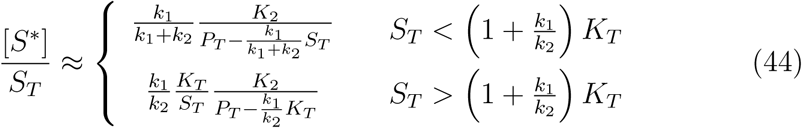

when *k*_1_*K_T_ < k*_2_*P_T_* (*α* < 1) and

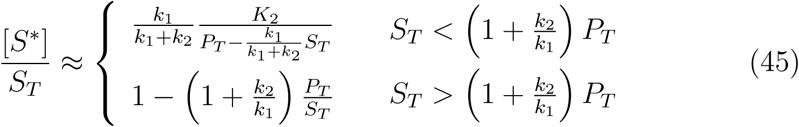

when *k*_1_*K_T_ > k*_2_*P_T_* (*α* > 1). Interestingly, when *α* < 1 the fraction of phosphorylated substrate changes in a nonmonotonic manner (Fig. 4B) with a maximum at (1 + *k*_1_/*k*_2_)*K_T_* (Eq. 44). In contrast, if *k*_1_*K_T_ > k*_2_*P_T_* the fraction of phosphorylated substrate remains low (since *K*_2_/*P_T_* ≪ 1) below the threshold and increases hyperbolically beyond that point (Eq. 45).

### 3.2. The high-K_M_ regime

To derive an equation for [*Y*\*] in the high-*K_M_* regime (*K*_1_ ≫ *K_T_* and *K*_2_ ≫ *P_T_*) we substitute the expressions from Eqs. (26) and (27) into Eq. (18) to obtain the approximation

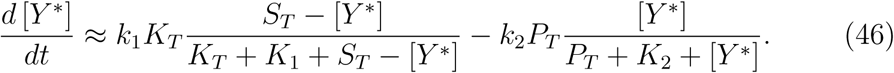

Hence, in the high-*K_M_* regime the dynamics of [*Y*\*] is described by a similar equation as [*S*\*] in the context of the Goldbeter-Koshland model (Eq. 10). The steady state solution for the total fraction of phosphorylated substrate is given by

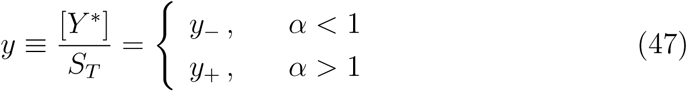

with

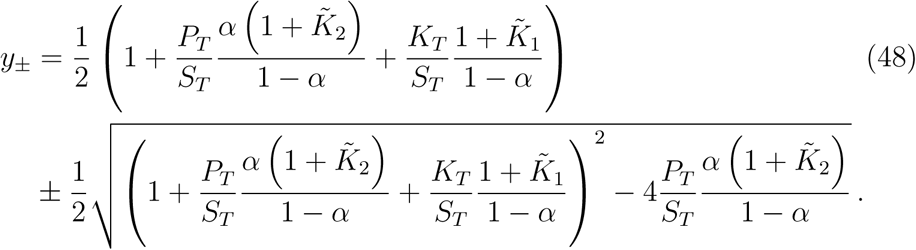

Here, 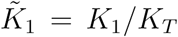 and 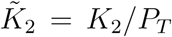 denote Michaelis-Menten constants which are scaled by enzyme (rather than substrate) concentrations. Note that the expression in Eq. (48) reduces to that of the Goldbeter-Koshland model (Eq. 12) when *S_T_* ≫ max(*K_T_, P_T_*) while keeping *K*_1_/*S_T_* and *K*_2_/*S_T_* constant.

### 3.2.1. Response curve with respect to k_1_

Under conditions when the substrate concentration is equal to or lower than that of the converter enzymes (*S_T_* ≤ min(*K_T_, P_T_*)) one may linearize the right-hand side of Eq. (46) to obtain the steady state approximation (Fig. 5A)

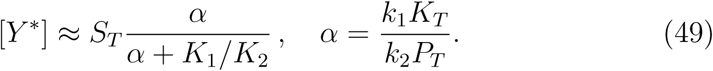

Here, [*Y*\*] increases in a hyperbolic manner as a function of *k*_1_ similar to [*S*\*] in the Goldbeter-Koshland model when both enzymes are unsaturated (Fig. 2B).

**Figure 5:**
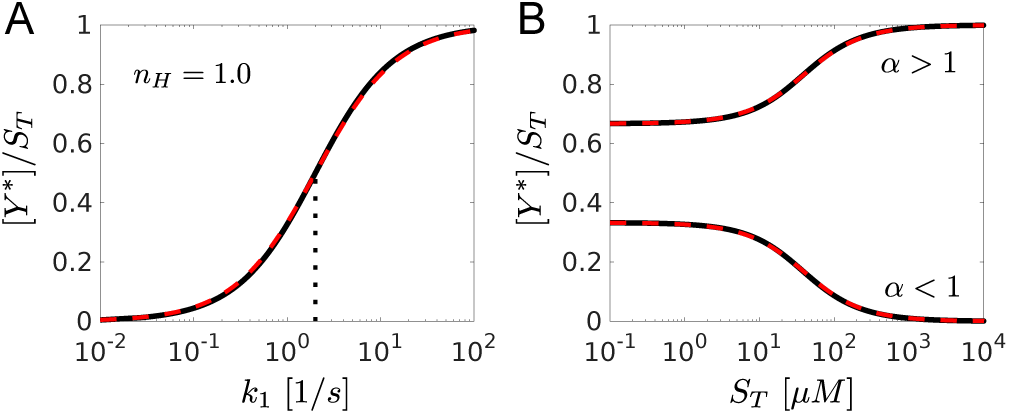
Response curves for the total fraction of phosphorylated substrate in the high-*K_M_* regime: (A) as a function of *k*_1_, (B) as a function of *S_T_* for different values of *α* = *k*_1_*K_T_*/(*k*_2_*P_T_*). Black solid lines correspond to the numerical steady state solution of Eqs. (1) - (6). Red dashed lines were computed from Eqs. (49) (A) and (47) (B). The dotted line indicates the value for half-maximal activation (A). Parameters: (A) *k*_2_ = 2/*s*, (B) *k*_2_ = 0.5/*s* (upper curve), *k*_2_ = 2/*s* (lower curve), *k*_1_ = 1/*s*, *K*_1_ = *K*_2_ = 10*μM*, *S_T_* = *K_T_* = *P_T_* = 1*μM*.

### 3.2.2. Response curve with respect to S_T_

When considered as a function of *S_T_* Eq. (49) is a poor approximation. In that case we employ the solution from Eqs. (47) and (48). Similar as in the low-*K_M_* regime the total fraction of phosphorylated substrate may decrease or increase depending on whether *α* < 1 or *α* > 1, respectively (Fig. 5B).

### 3.3. Mixed regime I: K_1_ ≪ K_T_ and K_2_ ≫ P_T_

To derive an equation for [*Y*\*] under conditions when the kinase is saturated while the phosphatase is unsaturated we substitute the expressions from Eqs. (27) and (28) into Eq. (18) to obtain the approximation

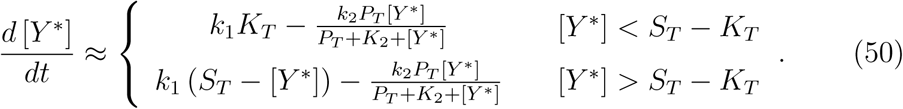

Setting the first line to zero yields

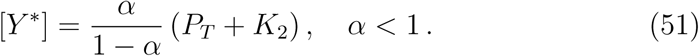

Since [*Y*\*] must be positive this solution only exists for *α* < 1. In the regime [*Y*\*] > *S_T_ − K_T_* the steady state concentration of *Y*\* is determined by the quadratic equation

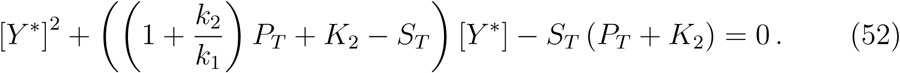

Since [*Y*\*] must be positive the solution of this equation reads

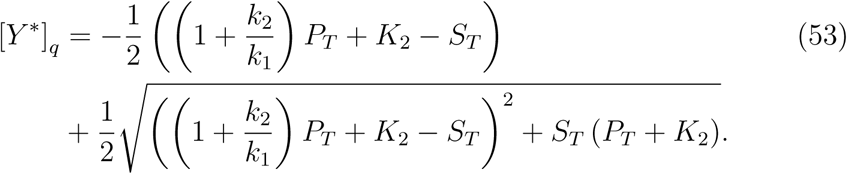

In the limit *P_T_ ≪ K*_2_ (which holds by assumption) and if *k*_2_/*k*_1_ ~ *O*(1) the solution of the quadratic equation (52) can be approximated by (cf. Appendix)

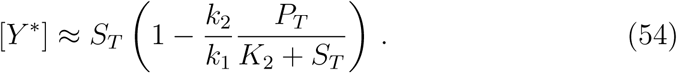

Hence, for *α* > 1 the steady state solution of Eq. (50) can be approximated by Eq. (53) (or by Eq. 54) whereas for *α* < 1 the approximation reads

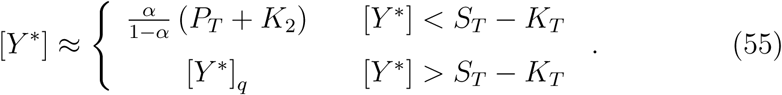

### 3.3.1. Response curve with respect to k_1_

As *k*_1_ increases from small to large values changes from the regime *α* < 1 to the regime *α* > 1 so that we have to combine the solutions in Eqs. (53) (or Eq. 54) and (55). To this end, we first compute the location of the threshold by solving

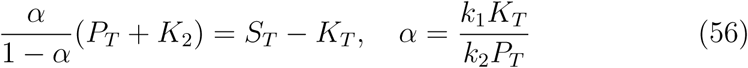

for *k*_1_ with the result

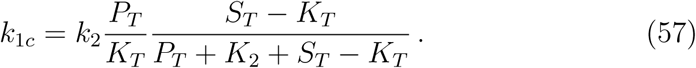

Note that this threshold is always smaller than that obtained from the condition *α* = 1. Also, the threshold only exists if *k*_1*c*_ > 0, i.e. if *K_T_ < S_T_*.

Hence, for *K_T_ < S_T_* we obtain from Eq. (55) the approximation (cf. Fig. 6A)

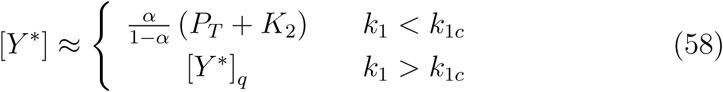

while for *K_T_ > S_T_* the approximation is given by Eq. (53) (or Eq. 54) (cf. Fig. 6B).

### 3.3.2. Response curve with respect to S_T_

For *α* > 1 the total fraction of phosphorylated substrate is given by Eq. (53) or, to leading order, by Eq. (54) as

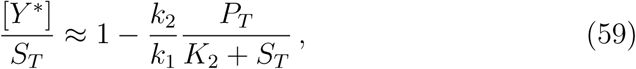

i.e. it increases monotonically as a function of *S_T_* (Fig. 6C).

In contrast, for *α* < 1 there exists a threshold defined by Eq. (56) so that Eq. (55) becomes

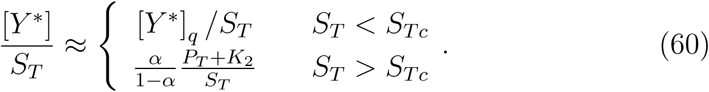

where *S_Tc_* is given by

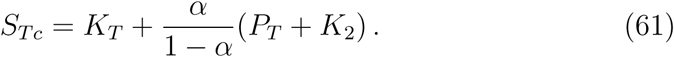

Hence, for *α* < 1 the total fraction of phosphorylated substrate exibits a maximum near *S_Tc_* (Eq. 61) since [*Y*\*]/*S_T_* is a decreasing function of *S_T_* beyond the threshold (Fig. 6D).

### 3.4. Mixed regime II: K_1_ ≫ K_T_ and K_2_ ≪ P_T_

To derive an equation for *Y*\* under conditions when the kinase is unsaturated while the phosphatase is saturated we substitute the expressions from Eqs. (26) and (29) into Eq. (18) to obtain the approximation

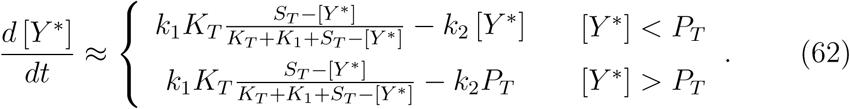

Setting the second line to zero yields

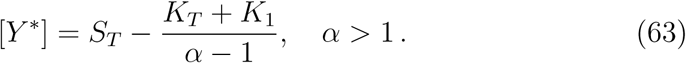

Since [*Y*\*] must be smaller than *S_T_* this solution only exists for *α* > 1. In the regime [*Y*\*] < *P_T_* the steady state concentration of *Y*\* is determined by the quadratic equation

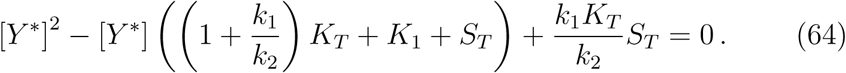

Since [*Y*\*] must not exceed *S_T_* the solution of this equation is given by

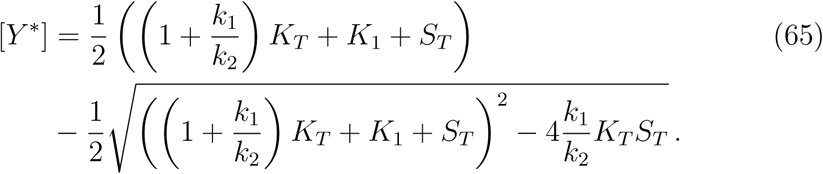

For 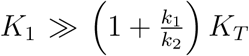 the solution of the quadratic equation (64) can be approximated by balancing the linear and the constant terms (Straube, 2014, 2015)

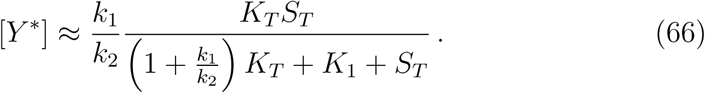

Combining these results yields for *α* > 1

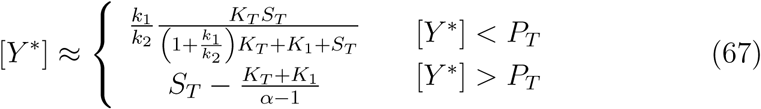

while for *α* < 1 the approximation is given by Eq. (66).

**Figure 6:**
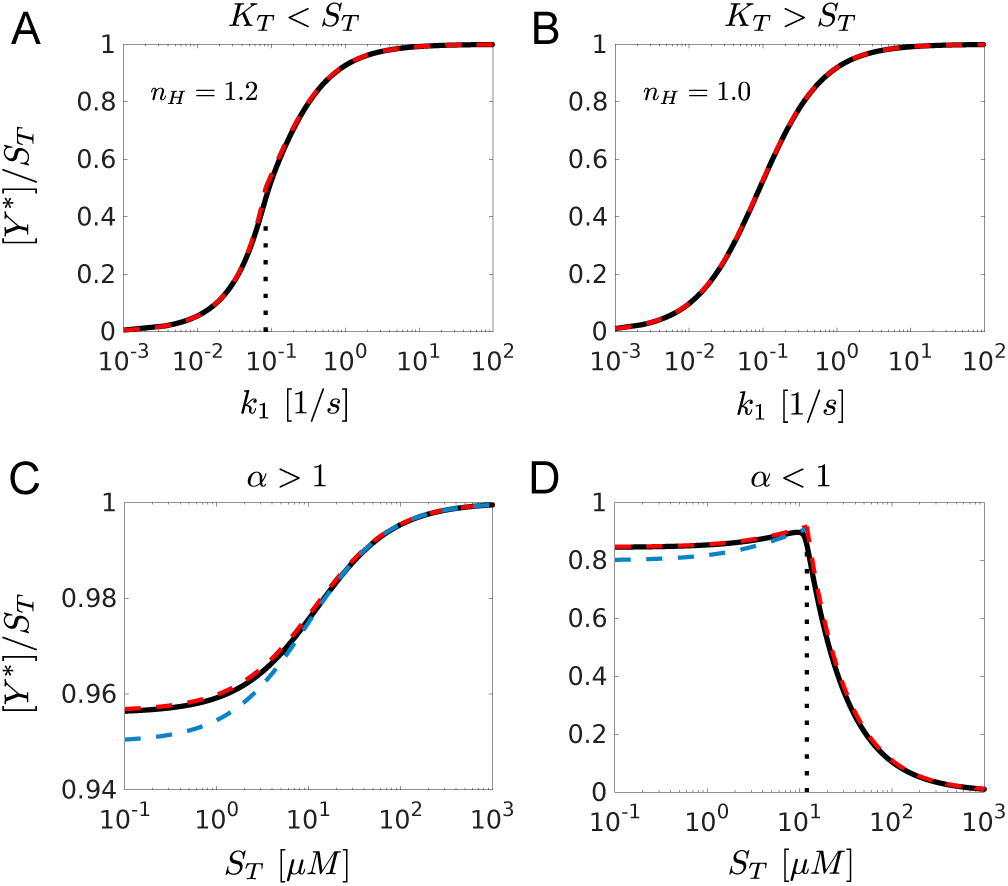
Response curves in the mixed regime I: *K*_1_ ≪ *K_T_* and *K*_2_ ≫ *P_T_*. (A,B) [*Y*\*] as a function of *k*_1_, (C,D) [*Y*\*]/*S_T_* as a function of *S_T_*. Black solid lines correspond to the numerical state solution of Eqs. (1) - (6). Red dashed lines were computed from Eq. (58) (A,B) and Eq. (60) (C,D) whereas blue dashed lines correspond to Eq. (59) (C,D). Dotted lines denote threshold values computed from Eq. (57) (A) and Eq. (61) (D). Parameters: *S_T_* = 2*μM*, *k*_2_ = 1/*s* (A), *S_T_* = 0.5*μM*, *k*_2_ = 1/*s* (B), *k*_2_ = 2/*s* (C) *k*_2_ = 0.5/*s* (D). Other parameters: *k*_1_ = 1/*s*, *K*_1_ = 0.01*μM*, *K*_2_ = 10*μM*, *K_T_* = *P_T_* = 1*μM*.

### 3.4.1. Response curve with respect to k_1_

As *k*_1_ increases from small to large values changes from the regime *α* < 1 to the regime *α* > 1 so that we have to combine the solutions in Eqs. (66) and (67). To this end, we first compute the location of the threshold by solving

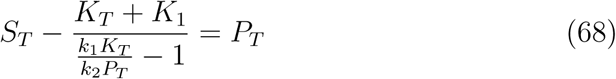

for *k*_1_ with the result

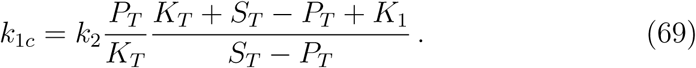

Note that this threshold is always larger than that obtained from the condition *α* = 1. Also, the treshold only exists (*k*_1*c*_ > 0) if *P_T_ < S_T_*. Hence for *P_T_ < S_T_* we obtain (Fig. 7A)

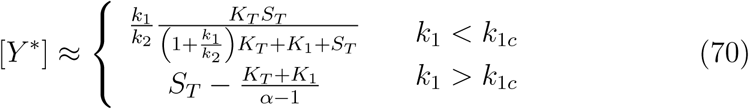

while for *P_T_ > S_T_* the approximation of the stimulus-response curve given by Eq. (66) (Fig. 7B).

### 3.4.2. Response curve with respect to S_T_

For *α* < 1 (and *k*_1_/*k*_2_ *~O*(1)) the total fraction of phosphorylated substrate can be approximated by (cf. Eq. 66)

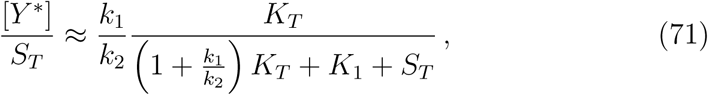

i.e. it decreases monotonically as a function of *S_T_* (Fig. 7C).

In contrast, for > 1 there exists a threshold defined by Eq. (68) so that Eq. (67) becomes

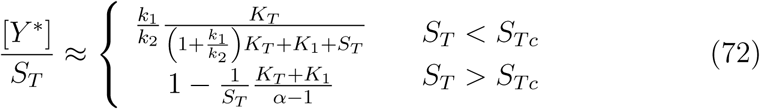

where *S_Tc_* is given by

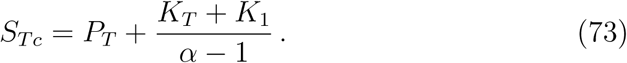

Hence, for *α* > 1 the total fraction of phosphorylated substrate exibits a minimum near *S_Tc_* (Eq. 73) since [*Y*\*]=*S_T_* is an increasing function of *S_T_* beyond the threshold (Fig. 7D).

**Figure 7:**
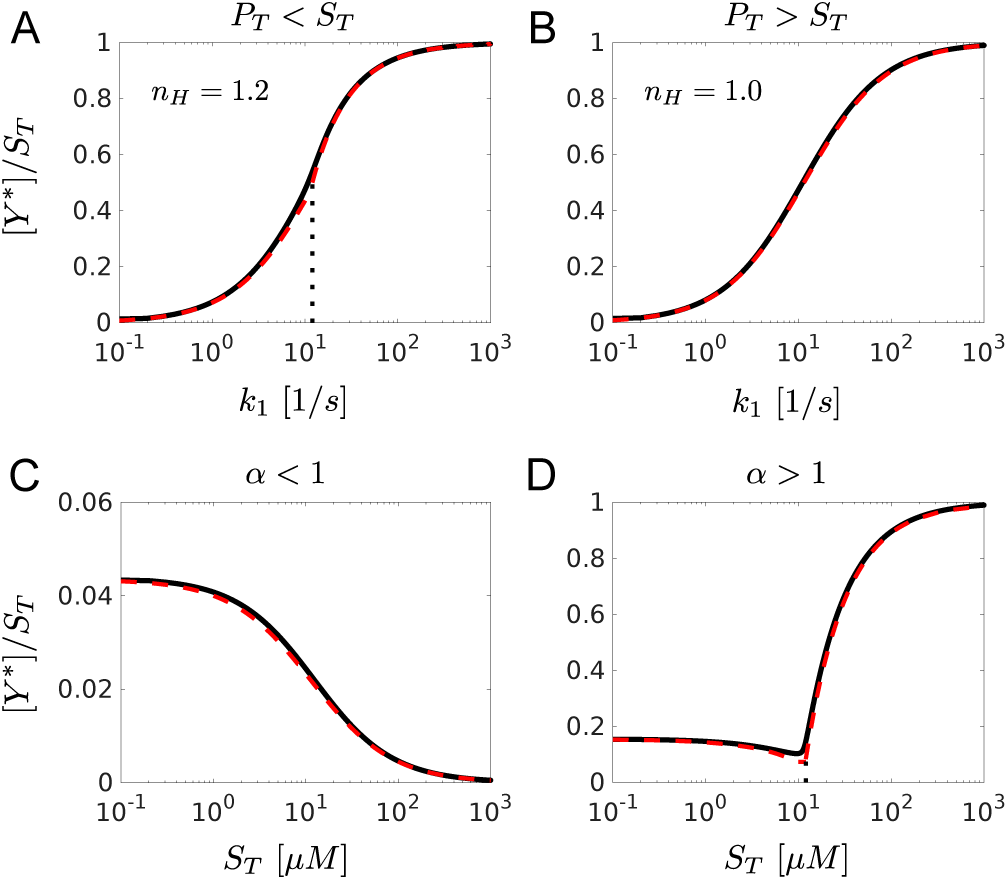
Response curves in the mixed regime II: *K*_1_ ≫ *K_T_* and *K*_2_ ≪ *P_T_*. (A,B) [*Y*\*] as a function of *k*_1_, (C,D) [*Y*\*]/*S_T_* as a function of *S_T_*. Black solid lines correspond to the numerical steady state solution of Eqs. (1) - (6). Red dashed lines were computed from Eq. (70) (A), Eq. (66) (B), Eq. (71) (C) and Eq. (72) (D). Dotted lines denote threshold values computed from Eq. (69) (A) and Eq. (73) (D). Parameters: *S_T_* = 2*μM*, *k*_2_ = 1/*s* (A), *S_T_* = 0.5*μM*, *k*_2_ = 1/*s* (B), *k*_2_ = 2/*s* (C) *k*_2_ = 0.5/*s* (D). Other parameters: *k*_1_ = 1/*s*, *K*_1_ = 10*μM*, *K*_2_ = 0.01*μM*, *K_T_* = *P_T_* = 1*μM*.

### 4. Discussion and Conclusion

In this study we have analyzed the steady state behavior of covalent modification cycles under conditions when the concentrations of the converter enzymes (*K* and *P*) are comparable to or larger than that of the substrate (*S*). To this end we have employed the total quasi-steady state approximation (tQSSA) which is based on the total concentration of the modified substrate ([*Y*\*] = [*S*\*] + [*S*P*]). From a theoretical point of view this procedure facilitates the analysis under conditions when enzyme concentrations are high because [*Y*\*] accounts for both free and enzyme-bound substrate forms. From a practical point of view the tQSSA has the advantage that [*Y*\*] (rather than [*S*\*]) is often the quantity that is accessible in experiments.

The tQSSA yields quadratic equations whose analysis suggests to classify the steady state behavior of [*Y*\*] based on the ratio between the enzyme’s Michaelis-Menten constants and the respective enzyme concentration. Since each of the two enzymes can be either saturated (*K_M_ ≪ E_T_*) or unsaturated (*K_M_ ≫ E_T_*), there are 4 steady state operating regimes as depicted in Fig. 8. Depending on the parameter of interest (with respect to which the response curve is to be computed) each regime can be subdivided into one or more subregimes. For example, if the total fraction of phosphorylated substrate ([*Y*\*]=*S_T_*) is the quantity of interest its behavior can be classified according to the ratio *α* = *k*_1_*K_T_*/(*k*_2_*P_T_*) between the maximal reaction rates of the converter enzymes. If *α* < 1 the total fraction of phosphorylated substrate decreases as a function of total substrate while the opposite is true for *α* > 1. Interestingly, when only one of the converter enzymes is saturated (mixed regimes) nonmonotonic behavior is possible where [*Y*\*]/*S_T_* either exhibits a maximum (*α* < 1, Fig. 6D) or a minimum (*α* > 1, Fig. 7D). This simple classification may aid the interpretation of overexpression data to decide which of the two enzyme activities dominates under a given condition.

One of the hallmarks of the Goldbeter-Koshland model is its prediction of ultrasensitivity (Goldbeter and Koshland Jr., 1981) for the free form of the modified substrate (*S*\*). However, its occurrence does not only require both converter enzymes to be saturated but also the substrate to be in excess (*S_T_* ≫ max(*K_T_, P_T_*)) – a condition that is rarely satisfied in living cells (Blüthgen et al., 2006; Legewie et al., 2008). In general, the shape of the stimulus-response curve for [*Y*\*] (as a function of the catalytic rate constant *k*_1_) depends on the concentration(s) of the saturated converter enzyme(s) (Fig. 8). For example, if both enzymes operate in saturation (low-*K_M_* regime) and the sum of the concentrations of both converter enzymes does not exceed the substrate concentration (*K_T_* + *P_T_ < S_T_*) ultrasensitive responses with *n_H_* ≫ 1 are possible (Fig. 3B) similar as in the Goldbeter-Koshland model. Our results suggest that enhanced sensitivity is still possible in the regime *K_T_* + *P_T_ > S_T_* if the concentration of at least one of the converter enzymes remains smaller than that of the substrate.

**Figure 8:**
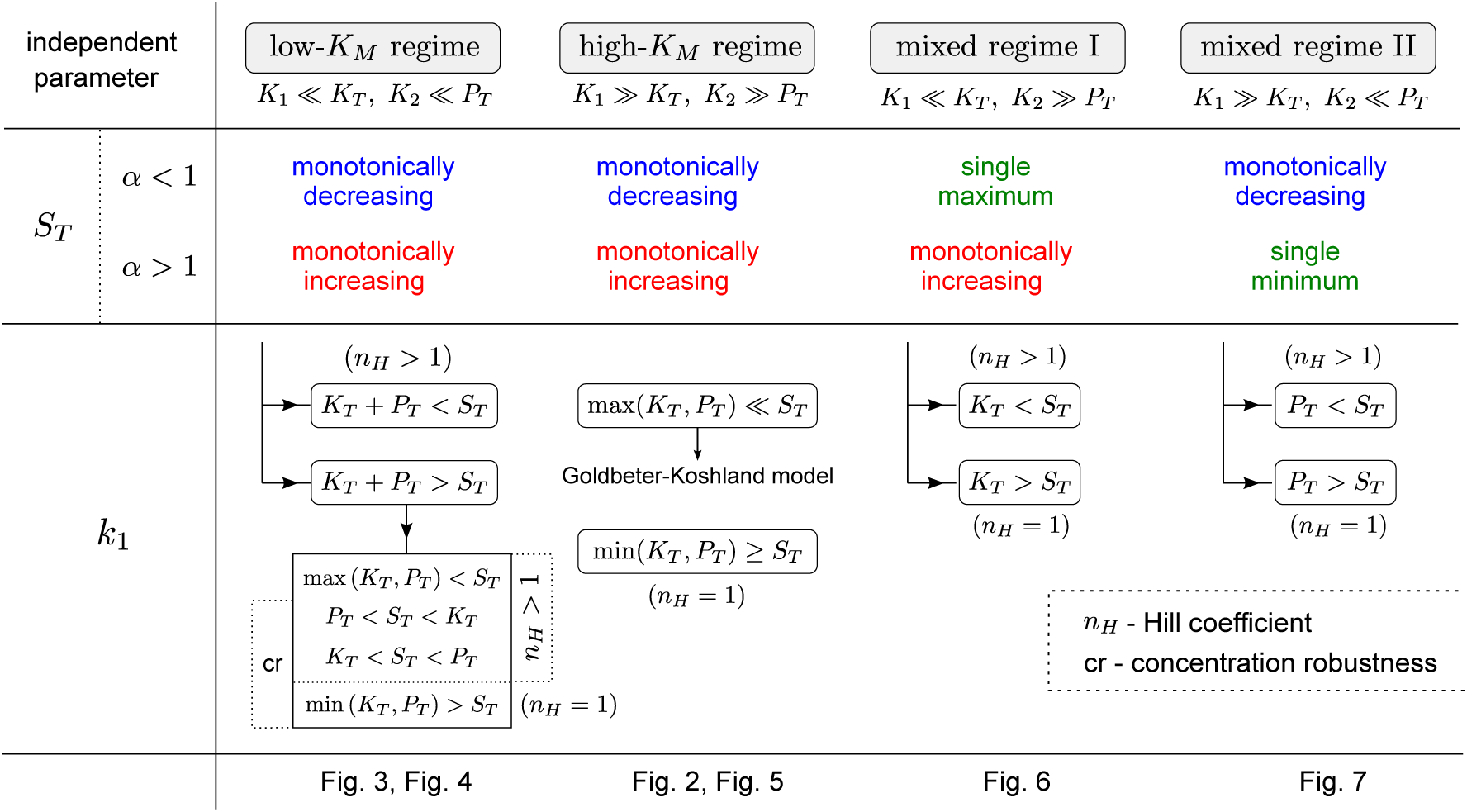
Classification of the steady state operating regimes of covalent modification cycles based on the tQSSA. 1st row: qualitative shape of the stimulus-response curve for [*Y*\*]/*S_T_* as a function of *S_T_* and *α* = *k*_1_*K_T_*/(*k*_2_*P_T_*). 2nd row: Classification of the stimulus-response curve for [*Y*\*] as a function of *k*_1_.

In the high-*K_M_* regime the steady state equation for [*Y*\*] becomes similar to that of the Goldbeter-Koshland model for [*S*\*]. This regime has been previously analyzed in the limit of substrate excess (*S_T_* ≫ max(*K_T_, P_T_*)) (Gomez-Uribe et al., 2007) where its predictions become identical to that of the Goldbeter-Koshland model. However, if the substrate concentration is lower than that of the converter enzymes the Hill coefficient of the response curve becomes *n_H_* = 1 (Fig. 5). In fact, we made similar observations in all 4 operating regimes (Fig. 8) suggesting that *S_T_* < min(*K_T_, P_T_*) is a sufficient condition for the loss of enhanced sensitivity at the level of [*Y*\*].

Concentration robustness is another network property that has received substantial attention from the theoretical side. The latter occurs if the stimulus-response curve is independent of the concentration of a protein in the network. In that case, changes in the expression level of that protein would not affect the steady state output of the system. In some cases concentration robustness is *absolute* in the sense that the steady state output of a system is completely independent of a protein concentration (Shinar and Feinberg, 2010). Often this form of robustness can be related to structural properties of the network such as bifunctionality of a converter enzyme (Shinar et al., 2007, 2009; Straube, 2013; Dexter et al., 2015). In our case, the concentration robustness is approximate in nature as it only occurs under certain conditions, i.e. if both converter enzymes are saturated and if the concentration of at least one of the converter enzymes is higher than that of the substrate. As a result concentration robustness can exist with respect *K_T_*, *P_T_* or both. Interestingly, the conditions for the occurrence of concentration robustness partially overlap with those for the occurrence of ultrasensitivity (cf. Fig. 8) suggesting that covalent modification cycles operating in the low-*K_M_* regime can simultaneously exhibit both concentration robustness and ultrasensitivity.

## Appendix A. Derivation of Eq. 54

Using the rescaled quantities

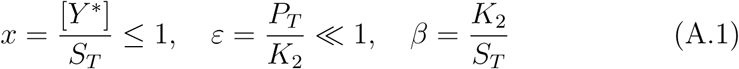

the quadratic equation (52) can be written in the form

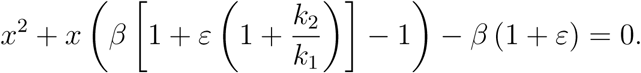

Assuming that *k*_2_/*k*_1_ *~ O* (1) we expand the solution as

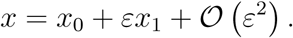

To lowest order this yields the quadratic equation

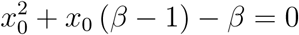

which has the two solutions

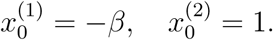

The next order correction is determined by

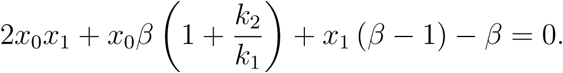

Substituting the positive solution 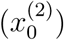 and solving for *x*_1_ yields

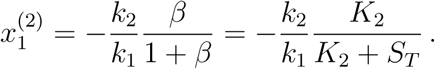

Together, this shows that

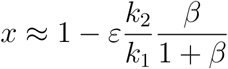

or (using Eq. A.1)

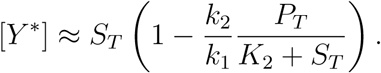

Note that this approximation remains positive in its range of validity since *P_T_ ≪ K*_2_ and *k*_2_/*k*_1_ *~ O* (1) by assumption.

